# Differential Modulation of Polycomb-Associated Histone Marks by cBAF, pBAF, and gBAF Complexes

**DOI:** 10.1101/2023.09.23.557848

**Authors:** Mary Bergwell, JinYoung Park, Jacob G. Kirkland

## Abstract

Chromatin regulators are a group of proteins that can alter the physical properties of chromatin to make it more or less permissive to transcription by modulating another protein’s access to a specific DNA sequence through changes in nucleosome occupancy or histone modifications at a particular locus. Mammalian SWI/SNF complexes (mSWI/SNF) are a group of ATPase-dependent chromatin remodelers that alter chromatin states. In mouse embryonic stem cells (mESCs), there are three primary forms of mSWI/SNF: canonical BAF (cBAF), polybromo-associated BAF (pBAF), and GLTSCR-associated BAF (gBAF or ncBAF). While cBAF and gBAF contain the SS18 protein subunit, pBAF lacks SS18. Previous studies used a novel dCas9-mediated inducible recruitment (FIRE-Cas9) of mSWI/SNF complexes via SS18 to the *Nkx2.9* locus. *Nkx2.9* is a developmentally regulated gene that requires mSWI/SNF for transcriptional activation during neural differentiation. However, in mESCs, *Nkx2.9* is bivalent, meaning nucleosomes at the locus have both active and polycomb-associated repressive modifications. Upon recruitment of SS18-containing complexes, polycomb-associated histone marks are removed, followed by transcriptional activation of *Nkx2.9*. However, since both cBAF and gBAF share the SS18 subunit, it is unclear whether one or both complexes oppose the polycomb repressive marks. The ability of pBAF to do the same also remains unknown. In this study, we used unique subunits to recruit each of the three complexes to the Nkx2.9 locus individually. Here, we show that cBAF most effectively opposes polycomb repressive marks at *Nkx2.9*, leading to transcriptional activation of the gene. Recruitment of cBAF complexes leads to a significant loss of the polycomb repressive-2 H3K27me3 and polycomb repressive-1 H2AK119ub histone marks, whereas gBAF and pBAF do not. Moreover, nucleosome occupancy alone cannot explain the loss of these marks. Our results demonstrate that cBAF has a unique role in the direct opposition of polycomb-associated histone modifications that gBAF and pBAF do not share.

1. Cell Cycle and Cancer Biology Program, Oklahoma Medical Research Foundation, Oklahoma City, OK, 73104
2. Department of Cell Biology, University of Oklahoma Health Science Center, Oklahoma City, OK, 73104

## Introduction

The eukaryotic genome is organized into three primary chromatin states: accessible euchromatin, conditionally accessible facultative heterochromatin, and mostly inaccessible constitutive heterochromatin. DNA accessibility is essential for regulating transcription factor binding; therefore, these accessibility states contribute to the transcriptional state, with the most accessible euchromatin being largely transcriptionally active and the two heterochromatin states being transcriptionally repressed. Facultative heterochromatin is generally defined by polycomb group (PcG) proteins and their respective histone marks. Polycomb consists of two main repressive complexes, polycomb repressive complex 1 (PRC1) and polycomb repressive complex 2 (PRC2), that modify histones as a part of transcriptional silencing. PRC1 writes and reads the H2AK119ub modification, while PRC2 writes and reads the H3K27me3 histone modification^1^. There is an additional cross-talk between these two complexes. Regulation of facultative heterochromatin—polycomb proteins and their respective histone modifications—is critical during development. For a cell to differentiate, it must coordinate new transcriptional programs^2–4^. At each point in the cell fate decision-making tree, new sets of genes must be activated while other sets of genes must be repressed. The mSWI/SNF family of protein complexes (also known as BAF complexes^5^) are chromatin proteins that help regulate the balance of facultative heterochromatin in the cell and are required to modulate transcriptional states during development^5–10^. In stem cells, there are three distinct, major forms of BAF complexes often in the same cell: canonical BAF (cBAF), Polybromo-associated BAF (pBAF), and GLTSCR-associated BAF (gBAF)^11–13^. The three forms share common subunits, but each complex has distinct subunits not shared by the others.

Although the earliest SWI/SNF studies described its role in nucleosome remodeling^14^, subsequent studies revealed its role in regulating chromatin accessibility and transcription through opposition to polycomb group proteins^7–10^. To understand how BAF complexes evict polycomb and activate genes, we developed a method of rapamycin-induced chromatin regulator recruitment to specific loci called FIRE-Cas9^15^. FIRE-Cas9 uses catalytically dead Cas9 and sgRNAs to target a particular locus. FRB and FKBP fusion proteins are then used to force chemical-induced proximity at the dCas9-bound locus by adding rapamycin to the cell culture media. The FIRE-Cas9 system enables the recruitment of a specific chromatin regulator and tracking of the resulting consequences with minute-by-minute kinetics on physiologic chromatin^15^. Our previous study used the BAF subunit SS18 as the recruitment anchor^15^. Though it was not known then, SS18 is present in both cBAF and gBAF complexes^12,13,16^. We found that recruiting SS18-containing complexes led to the rapid eviction of polycomb proteins, and this preceded transcriptional activation, thus elucidating the order of events of transcriptional regulation on the chromatin level by BAF complexes^8,15^. However, since both cBAF and gBAF share the SS18 subunit, it is unclear whether one or both complexes oppose the polycomb repressive marks. The ability of the third major complex, pBAF, to do the same also remains unknown.

In this study, we describe a modification of the FIRE-Cas9 system using tagged subunits that are found in only one of the three complexes as recruitment anchors. By integrating the unique subunits DPF2 (cBAF), BRD9 (gBAF), and PHF10 (pBAF) into our FIRE-Cas9 system, we can specifically recruit cBAF, gBAF, or pBAF. With this approach, we can perform experiments that will define and distinguish the contributions of the three complexes to polycomb-associated histone modifications at the bivalent *Nkx2.9* gene. The recruitment studies presented here show differential modulation of polycomb-associated histone marks by cBAF, gBAF, and pBAF complexes.

## Results

### Tagged unique BAF subunits are expressed and incorporated into BAF complexes

To achieve better fusion protein expression, we modified the FIRE-Cas9 system by switching the Frb/Fkbp dimerization tags. Previously, the Fkbp tag was attached to the MS2 bacteriophage coat protein (MS2), and the Frb tag was attached to our chromatin regulator of interest. In this study, we have now tagged MS2 with Frb (MS2-2xFrb) and our chromatin regulator with Fkbp domains (CR-2xFkbp-V5) (Figure 1A). We used lentiviral transduction to express each of the FIRE-Cas9 components in mouse embryonic stem cells (mESCs) (Figure 1B). Our experimental lines were built by first transducing with expression plasmids for dCas9-HA and MS2-2xFrb, ensuring equal expression in the final lines (Figure 1C). We then transduced cells with DPF2-2xFkbp-V5, BRD9-2xFkbp-V5, or PHF10-2xFkbp-V5 (Figure 1D). We finally transduced these lines with three sgRNA with extra stem-loop structures targeting the promoter of *Nkx2.9* to complete the system. The sgRNAs guide the dCas9 to the *Nkx2.9* promoter, and the MS2-Frb protein binds the stem-loops. Only after adding rapamycin are specific BAF complexes recruited to *Nkx2.9* by the dimerization of the Frb and Fkbp domains (Figure 1A). dCas9 is always present at the locus in our control (no rapamycin) and experimental conditions (rapamycin), allowing for better comparisons than methods where a chromatin regulator is directly tethered to dCas9. A key advantage of chemical-induced proximity^17^ is that the rapamycin selectively binds to both Frb and Fkbp at once with an affinity 2000-fold tighter than binding to the Frb domain (found in mTOR) alone, which allows us to use rapamycin at low doses (3 nM) in FIRE-Cas9 experiments^18^. We performed a series of co-immunoprecipitations to ensure that our tagged proteins were incorporated into BAF complexes. First, we immunoprecipitated with a mouse anti-V5 antibody to pull down our CR-2xFkbp-V5 fusion proteins and then performed western blots to other shared BAF subunits BAF155 (SMARCC1) and BRG1 (SMARCA4). BAF155 forms a dimer as the initial BAF core in the first step of BAF complex assembly^19^. Only after this core is formed does the assembly process diverge for the three major complexes^19^. The unique subunits tagged in this study are subsequently added, creating the cBAF, gBAF, and pBAF cores^19^. Finally, the ATPase module, which includes BRG1, is added in one of the final steps of complex assembly^19^. Here, we show that pulldown using an antibody to the V5 epitope, DPF2-2xFkbp-V5, BRD9-2xFkbp-V5, and PHF10-2xFkbp-V5 all interact with BAF155 and BRG1, suggesting they are part of full BAF complexes (Figure 1E). To further confirm PHF10-2xFkbp-V5 interactions with pBAF-specific subunits, we performed a pulldown with ARID2. These data show that ARID2 interacts with the tagged PHF10-2xFkbp-V5 subunit (detected with a V5 antibody), along with BAF155 and BRG1 (Figure 1F). Detection of PHF10-2xFkbp-V5 was specific to our expressing line and was not found in wild-type TC1 cells. Finally, we pulled down using an antibody to PHF10 itself and found the V5-tagged version of PHF10 in the pulldown interacting with BAF155 and BRG1 (Supplemental Figure 1). These data suggest that DPF2-2xFkbp-V5, BRD9-2xFkbp-V5, and PHF10-2xFkbp-V5 are all incorporated into their respective BAF complexes.

**Figure 1.**
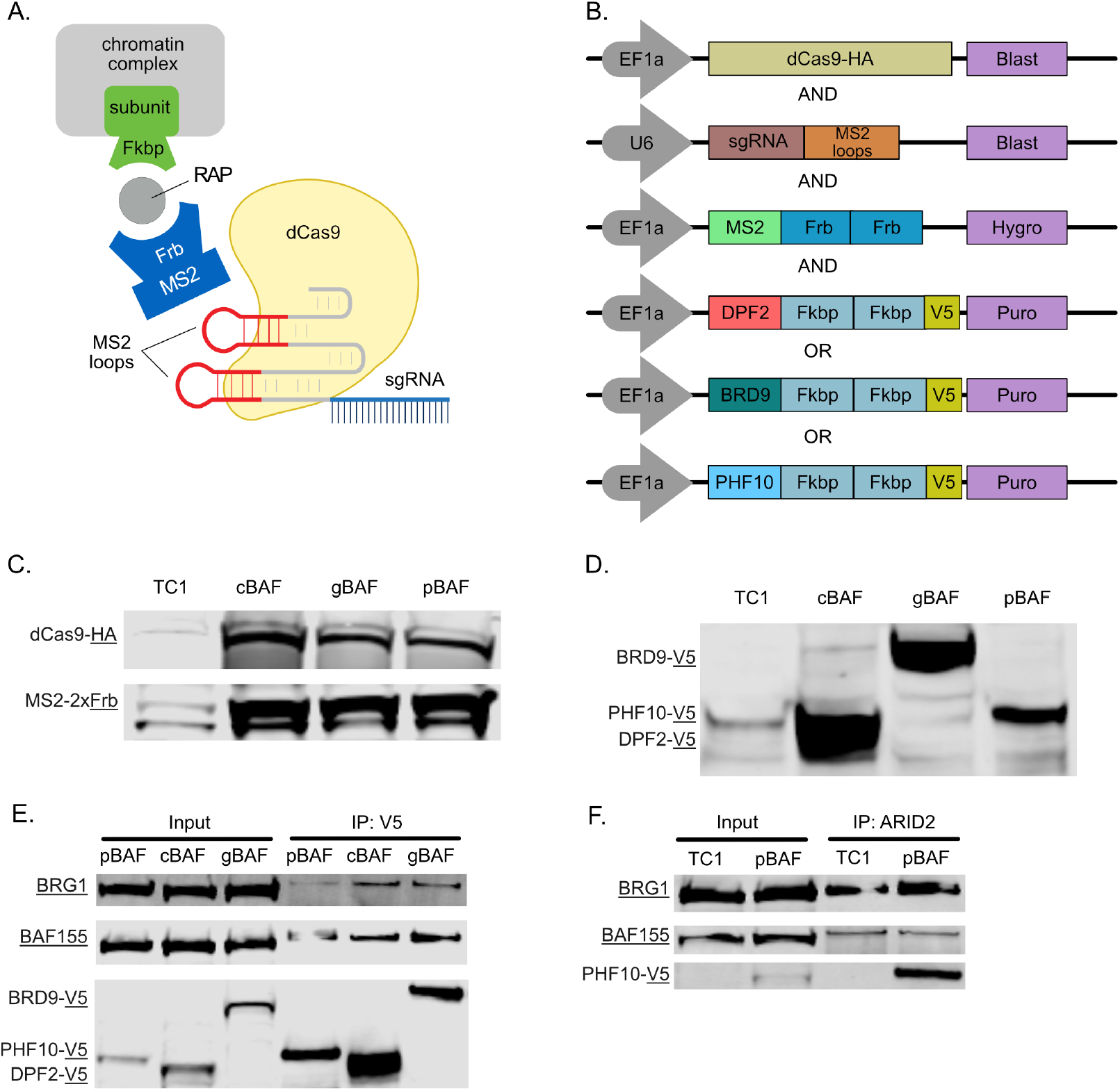
Tagged unique BAF subunits are expressed and incorporated into BAF complexes. **A**. Schematic of the modified FIRE-Cas9 system showing induced proximity by dimerization of Frb and Fkbp domains by rapamycin. **B**. Lentiviral expression constructs of FIRE-Cas9 components. **C**. Western blot showing expression of dCas9-HA and MS2-2xFrb in each cell line with wild-type TC1 negative control. **D**. Western blot showing expression of 2xFkbp-V5 tagged BAF subunits used for recruitment of their respective complexes. **E**. Co-IP pulldown with an antibody against V5 followed by Western blot showing interactions between V5 tagged BAF subunits and the core subunit BAF155 and the ATPase BRG1. **F**. Co-IP pulldown with an antibody against the pBAF specific subunit ARID2 showing interactions with BRG1, BAF155, and PHF10-2xFkbp-V5.

### Recruitment of specific BAF complexes via unique subunits

After showing that our tagged DPF2, BRD9, and PHF10 subunits are incorporated into complete BAF complexes, we recruited cBAF, gBAF, or pBAF to the *Nkx2.9* promoter. Here, we added rapamycin for 24 hours to achieve prolonged chromatin regulation by BAF complexes, followed by fixation and chromatin immunoprecipitation (ChIP) and qPCR to evaluate recruitment-based changes in polycomb-associated histone marks. To show that we are recruiting our tagged subunits, we performed ChIPs using a V5 antibody since each of our recruitment subunits contains this epitope followed by qPCR (Figure 2A-B). DPF2-based cBAF recruitment showed a 20-fold enrichment, while BRD9-based gBAF recruitment was 5.4-fold (Figure 2A-B). However, PHF10-based pBAF recruitment failed to show enrichment via the V5 antibody (Supplemental Figure 2A). We then performed ChIP against the PHF10 subunit itself and found robust enrichment of the recruited subunit (7.8-fold Figure 2C). This observation complicates our ability to directly compare the overall recruitment levels of the different complexes since even the same antibody shows a differential ability to bind the epitope (V5 in this case) depending on the subunit tagged and the complex it incorporates into. However, in each case, enrichment of the recruited subunit is specifically around the sgRNA binding sites (spanning −172 to −25bp) and does not spread far up or downstream.

**Figure 2.**
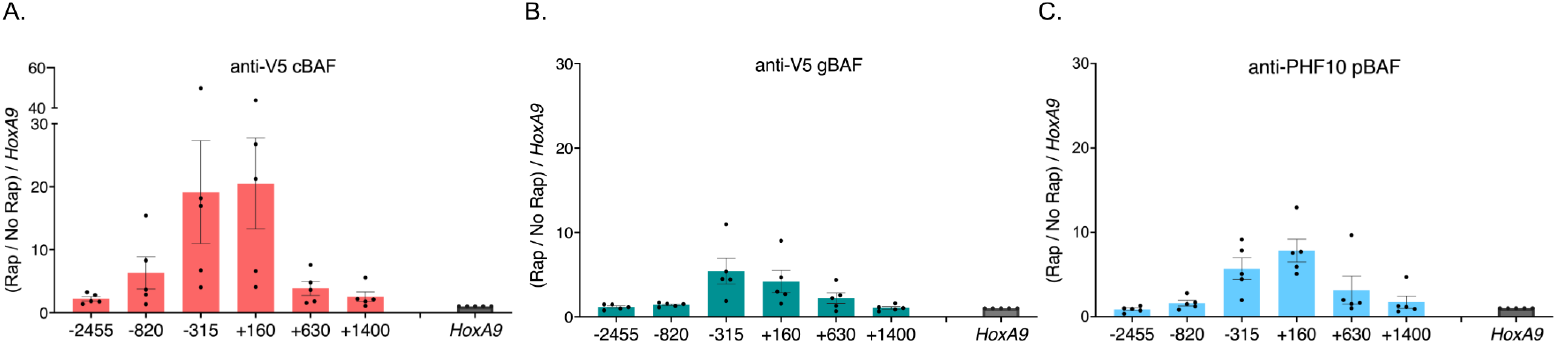
Recruitment of specific BAF complexes via unique subunits. Chromatin immunoprecipitations followed by qPCR detecting enrichment of the unique recruited subunits. 0 bp denotes the TSS of *Nkx2.9* and the recruitment site spans −172 to −25bp (upstream of the TSS). Data is presented as a fold change of recruitment/no recruitment normalized to *HoxA9*. **A**. Pulldown of DPF2-2xFkbp-V5 with an antibody against V5 (cBAF). **B**. Pulldown of BRD9-2xFkbp-V5 with an antibody against V5 (gBAF). **C**. Pulldown of PHF10-2xFkbp-V5 with an antibody against V5 (pBAF). (n=5; s.e.m.)

### Only cBAF complexes robustly oppose polycomb-associated histone marks

Next, we evaluated polycomb-associated histone marks. H3K27me3 is associated with PRC2 complexes, while H2AK119ub is associated with PRC1 complexes^1^. Here, we found critical differences between the BAF complexes. Our previous work showed that SS18-containing complexes led to the loss of repressive H3K27me3 histones upon recruitment to *Nkx2.9*^*15*^. However, since both cBAF and gBAF contain SS18, it remained unclear whether both or only one of these complexes was responsible for this observation. We now show that cBAF complexes are predominantly responsible for H3K27me3 loss with a minor gBAF-mediated loss (Figure 3A-B). Like gBAF, pBAF complexes only provide minimal loss of the H3K27me3 mark and only directly at the recruitment site, failing to spread like cBAF-mediated recruitment (Figure 3C). All three complexes evicted H3K27me3 better than H2AK119ub, including cBAF complexes (Figure 3D-F). Once again, cBAF was the most robust evictor of PRC1-associated marks (Figure 3D). Additionally, eviction of H2AK119ub occurred most strongly at the recruitment site and spread upstream of the promoter but not downstream of the promoter in the gene body in contrast with H3K27me3 (Figure 3D-F).

**Figure 3.**
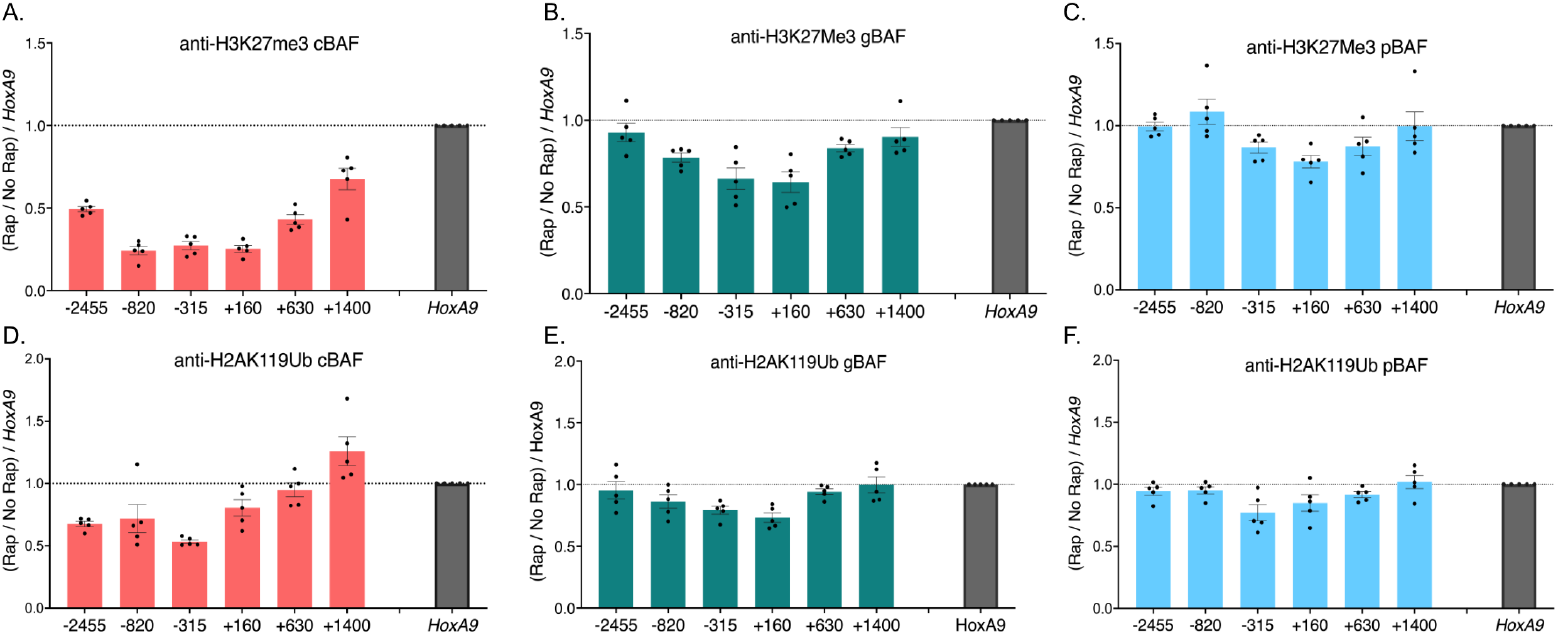
Only cBAF complexes robustly oppose polycomb-associated histone marks. Chromatin immunoprecipitations followed by qPCR detecting enrichment of histone modifications. 0 bp denotes the TSS of *Nkx2.9* and the recruitment site spans −172 to −25bp (upstream of the TSS). Data is presented as a fold change of recruitment/no recruitment normalized to *HoxA9*. **A**. Pulldown of H3K27me3 after recruitment of cBAF complexes **B**. Pulldown of H3K27me3 after recruitment of gBAF complexes **C**. Pulldown of H3K27me3 after recruitment of pBAF complexes **D**. Pulldown of H2AK119ub after recruitment of cBAF complexes **E**. Pulldown of H2AK119ub after recruitment of gBAF complexes **F**. Pulldown of H2AK119ub after recruitment of pBAF complexes (n=5; s.e.m.)

### Nucleosome depletion is insufficient to explain cBAF-mediated loss of polycomb-associated histone marks

Loss of a histone mark can occur through several mechanisms: (1) Co-recruitment of an enzyme or enzymes that remove the marks (demethylases and deubiquitinases in this case), leaving the rest of the nucleosomes intact; (2) Removal of a nucleosome, leaving a nucleosome-depleted region (NDR); or (3) Removal of a nucleosome which is then replaced with an unmodified nucleosome. To test these possibilities, we performed recruitment experiments of the three BAF complexes followed by ChIP using an antibody to Histone H4 (Figure 4A-C) or the C-terminus of H3, which detects nucleosomes regardless of H3 modifications (Figure 4D-F). We found that while there was some nucleosome loss following cBAF recruitment (Figure 4A and D), the loss of modified histones was larger than the loss of all nucleosomes, suggesting that model (2) removal of a nucleosome, leaving an NDR is insufficient to explain the loss of H3K27me3 (Figure 4G). H2AK119ub normalized to H3 nucleosome occupancy showed a more peculiar pattern where loss upstream of the marked histone was greater than that of all nucleosomes (Figure 4J). H2AK119ub downstream of the transcription starting site (TSS) showed the opposite pattern, where NDR explained all the losses of H2AK119ub. In fact, there was a slight increase in the amount of H2AK119ub per nucleosome downstream of the TSS. In contrast, gBAF recruitment led to minimal to no nucleosome depletion (Figure 4B and E), and any loss of H3K27me3 or H2AK119ub can be explained by nucleosome depletion (Figure 4H and K). Finally, pBAF recruitment led to minimal to no loss of H4 or H3 (Figure 4C and F), and the ratios of H3K27me3 and H2AK119ub on a per nucleosome level remained unchanged (Figure 4I and L).

**Figure 4.**
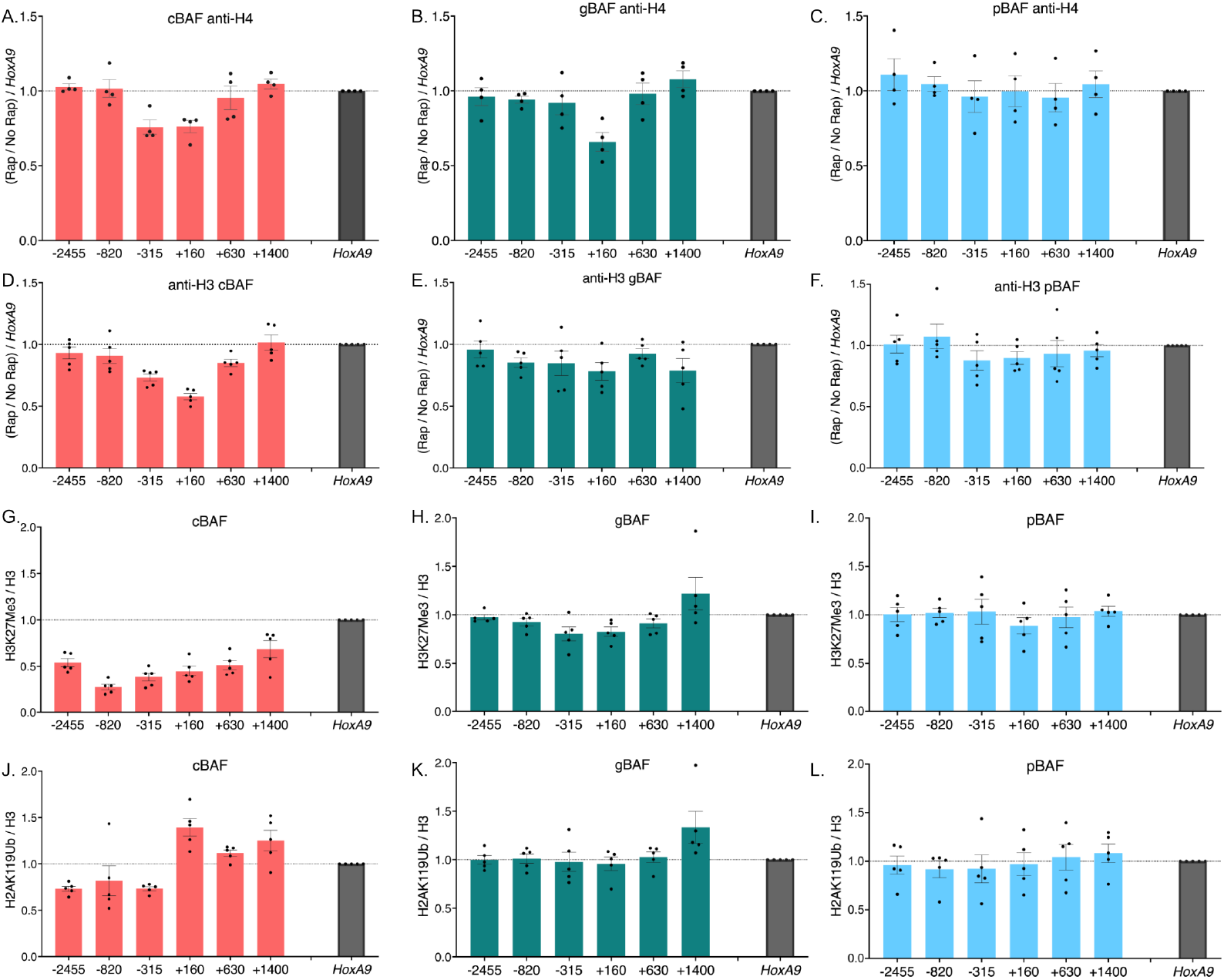
Nucleosome depletion is insufficient to explain cBAF-mediated loss of polycomb-associated histone marks. Chromatin immunoprecipitations followed by qPCR detecting enrichment of histones and their modifications. The recruitment site is approximately −100bp. 0 bp denotes the TSS of *Nkx2.9*. Data is presented as a fold change of recruitment/no recruitment normalized to *HoxA9*. **A**. Pulldown of H4 after recruitment of cBAF complexes **B**. Pulldown of H4 after recruitment of gBAF complexes **C**. Pulldown of H4 after recruitment of pBAF complexes **D**. Pulldown of H3 after recruitment of cBAF complexes **E**. Pulldown of H3 after recruitment of gBAF complexes **F**. Pulldown of H3 after recruitment of pBAF complexes **G**. The ratio of H3K27me3 over H3 after recruitment of cBAF complexes **H**. The ratio of H3K27me3 over H3 after recruitment of gBAF complexes **I**. The ratio of H3K27me3 over H3 after recruitment of pBAF complexes **J**. The ratio of H2AK119ub over H3 after recruitment of cBAF complexes **K**. The ratio of H2AK119ub over H3 after recruitment of gBAF complexes **L**. The ratio of H2AK119ub over H3 after recruitment of pBAF complexes (n=4-5; s.e.m.)

### cBAF and gBAF recruitment leads to a gain of H3.3 containing nucleosomes

Since we found that nucleosome depletion was insufficient to explain the loss of polycomb-associated histone marks, we set out to test model (3), whereby the removal of a polycomb-associated nucleosome is then followed by the replacement with an unmodified nucleosome. Outside of the centromere, mammalian nucleosomes may contain one of 3 Histone 3 variants: H3.1/H3.2 (replication-dependent deposition) or H3.3 (replication-independent deposition). Here, we hypothesized that the replacement nucleosomes may be deposited independently of replication. H3.3 differs from H3.1 and H3.2 by 4 or 5 amino acids, so we used an antibody raised against this variable region and is specific to the H3.3 variant. Here, we found an increase in H3.3 histones after the recruitment of cBAF (Figure 5A). Curiously, we also saw an increase in H3.3 even though we didn’t see a large loss of polycomb-associated histone marks after the recruitment of gBAF (Figure 3B and E). There was no gain in H3.3 after the recruitment of pBAF complexes. These data contrast the data when we used an antibody that recognizes all of H3.1, H3.2, and H3.3, whereby levels of H3 were depleted or remained unchanged (Figure 4). Therefore, these data are consistent with a model whereby cBAF evicts nucleosomes with polycomb-associated modifications, which are then replaced with H3.3 containing nucleosomes that largely lack H3K27me3 modifications.

**Figure 5.**
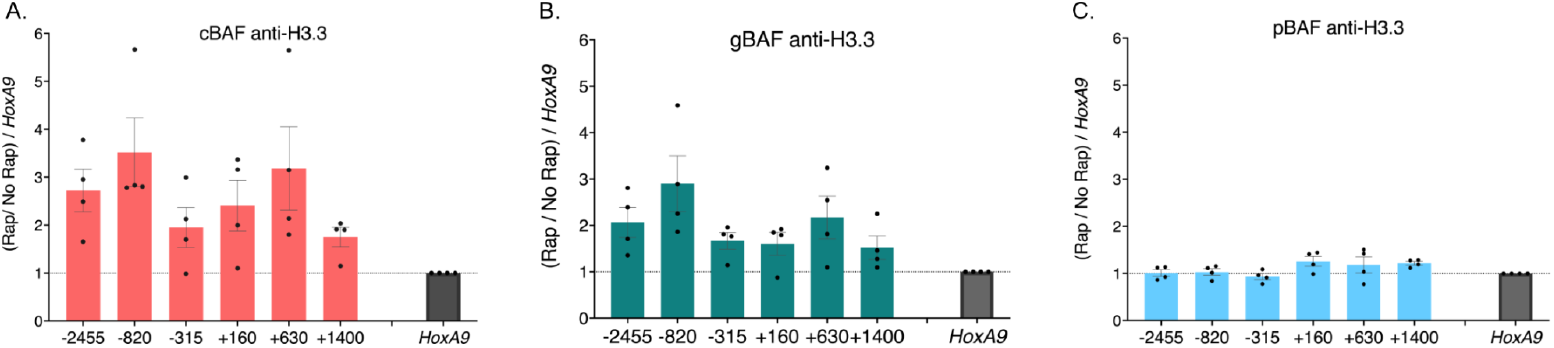
Recruitment of cBAF and gBAF complexes lead to a gain in histone variant H3.3. Chromatin immunoprecipitations followed by qPCR detecting enrichment of histones and their modifications. The recruitment site is approximately −100bp. 0 bp denotes the TSS of *Nkx2.9*. Data is presented as a fold change of recruitment/no recruitment normalized to *HoxA9*. **A**. Pulldown of H3.3 after recruitment of cBAF complexes **B**. Pulldown of H3.3 after recruitment of gBAF complexes **C**. Pulldown of H3.3 after recruitment of pBAF complexes (n=4 s.e.m.)

### cBAF is a transcriptional activator of Nkx2.9

Finally, we performed RT-qPCR to evaluate the ability of the different BAF complexes to promote transcriptional activation of *Nkx2.9*. We added rapamycin to the media for 24 hours to recruit the various BAF complexes. cBAF complexes acted as the most robust transcriptional activators, achieving a 2.8-fold increase in *Nkx2.9* mRNA. Compared to cBAF, neither gBAF (1.69-fold) nor pBAF (1.58-fold) were as strong of transcriptional activators of *Nkx2.9*, consistent with their inability to effectively evict polycomb-associated histone marks (Figure 6; p=0.039 cBAF vs. gBAF two-tailed t-test; p=0.0020 cBAF v.s pBAF two-tailed t-test). Because the fold change in transcripts appeared modest, we used CRISPR-A^20^ as a positive control. In this case, the activating Cas9 is constitutively present at the *Nkx2.9* as this system is not inducible like FIRE-Cas9, so gene activation has been happening for much longer than 24 hours. Therefore, CRISPR-A is expected to be a more potent transcriptional activator but represents a maximum activation limit. In these experiments, we compared a line expressing CRISPR-A without targeting sgRNAs to one expressing CRISPR-A with the same sgRNAs used to recruit the FIRE-Cas9 to *Nkx2.9*. We found that cBAF activated transcription of *Nkx2.9* to a level about half as strong as the CRISPR-A positive control (Supplemental Figure 4; p=0.17 Welch’s t-test).

**Figure 6.**
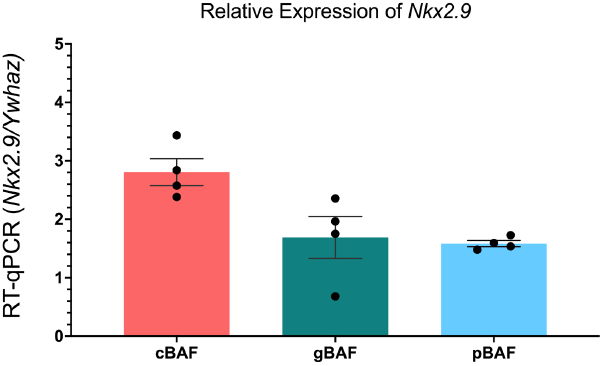
cBAF is a transcriptional activator of Nkx2.9. RT-qPCR of the fold-change of *Nkx2.9* transcription (normalized to *Ywhaz*) after recruitment of cBAF, gBAF, and pBAF. (n=4; s.e.m.)

## Discussion

We previously showed that BAF complexes that contain the SS18 subunit can evict H3K27me3 and activate transcription at the *Nkx2.9* gene. At the time, SS18 had only been described as a subunit of the BAF complex, but subsequent studies separated two SS18-containing complexes: the canonical BAF (cBAF) and GLTSCR-associated BAF (gBAF) complexes. Here, we show that we can specifically recruit cBAF, gBAF, and pBAF complexes through their unique subunits DPF2, BRD9, and PHF10 using our FIRE-Cas9 inducible recruitment system. We show that of the two SS18-containing complexes, cBAF is the more potent H3K27me3 (PRC2) and H2AK119ub (PRC1) opposing BAF complex, definitively showing a separation of the function of the two SS18-containing complexes, thus resolving ambiguity introduced by the discovery of gBAF complexes.

Loss of a histone mark can occur through several mechanisms: (1) Co-recruitment of an enzyme or enzymes that remove the marks, leaving the rest of the nucleosomes intact; (2) Removal of a nucleosome, leaving a nucleosome-depleted region (NDR); or (3) Removal of a nucleosome which is then replaced with an unmodified nucleosome. Our data show that the cBAF-mediated loss of these marks, particularly H3K27me3, cannot be explained by nucleosome depletion alone (model 2), as more modified histones are removed following cBAF recruitment than total histones. While future work is required to distinguish between models 1 and 3, we currently have no data consistent with model 1. For model 1 to be correct, cBAF must recruit KDM6A or KDM6B^21^ to remove H3K27me3 marks and BAP1/ASXL1^22,23^ to remove H2AK119ub marks at the same sites. This co-recruitment seems unlikely compared to model 3, where cBAF removes polycomb-associated nucleosomes, thereby removing H3K27me3 and H2AK119ub simultaneously. Following the removal of these nucleosomes, new nucleosomes containing H3.3 without H3K27me3 are inserted into the chromatin, which cBAF complexes may have a lower affinity for removing again, compared to polycomb-associated H3.1 containing nucleosomes. This specificity may be through interactions with PRC1 proteins themselves^8,9^, which are bound at these modified nucleosomes and aren’t found at the unmodified nucleosomes, thereby instructing cBAF which nucleosomes are to be preferentially evicted. However, our observation that H3K27me3 is removed more efficiently and over a broader region than H2AK119ub suggests additional undiscovered mechanisms of PRC2 eviction. cBAF may also preferentially avoid evicting H3.3-containing nucleosomes regardless of the polycomb-associated histone modifications. The BAF subunits BRG1 and SMARCB1 interact with the H3.3 chaperone HIRA complex^24^. This interaction may direct or promote H3.3 deposition at BAF-regulated sites, which then promotes transcription and a more open chromatin state^25^. However, the BRG1 and SMARCB1 subunits are shared between all BAF complexes, so it remains unclear how different BAF complexes influence H3.3 deposition to varying degrees.

We further show differential modulation of polycomb-associated histone marks by investigating pBAF complexes (which don’t contain SS18) for the first time. Here, we show that pBAF complexes are similar to the gBAF complex in their inability to oppose polycomb-associated histone marks at Nkx2.9 efficiently. Finally, when recruited to a bivalent gene, we show that the cBAF complex is a better transcriptional activator than the gBAF and pBAF complexes. In total, we have provided data describing different facultative heterochromatin modulations by the three main BAF complexes in mESCs. This suggests that each BAF complex has a unique role in changing or maintaining the balance of facultative heterochromatin in the nucleus when brought to a locus through native protein-protein interactions, which may include histone modifications, transcription factors, and other chromatin regulators.

### Limitations

One limitation of this study is that it proved impossible to measure the recruitment rates of the three BAF complexes in relation to each other. While all complexes contained a V5-tagged subunit, using a V5 antibody in the PHF10-2xFkbp-V5-based pBAF recruitment experiments showed no enrichment, whereas an antibody against PHF10 did. This suggests the V5 antibody had variable accessibility to its epitope within a formaldehyde-fixed BAF complex. We hypothesize that the V5 epitope is masked within a pBAF complex by other subunits after fixation since the epitope can be detected in denatured samples by western blot and pulled down in unfixed co-IPs. This also suggests that other subunits in the complexes have different epitope availability and show different cross-linking efficiencies for the same subunit. Our previous study used a custom-made rabbit polyclonal antibody to BAF155 to show the recruitment of another subunit within the SS18-containing complexes^15^. Unfortunately, the quantities of this antibody needed for this study are no longer available, nor did we previously test its ability to detect pBAF recruitment. We attempted ChIPs using commercially available antibodies to BAF155 and BRG1, present in all mESC BAF complexes, without success. So, we could only show the enrichment of other subunits not found in all BAF complexes, such as BAF57, ARID1A, and SS18, making comparisons of the relative enrichment of the different BAF complexes difficult.

## Materials and Methods

### Cell Culture

HEK293T cells were obtained from ATCC (CRL-3216) and cultured using standard parameters using DMEM media (Gibco 11960044) containing 10% FBS (Omega), 1% Glutamax (Gibco 35050061), and 1% Penicillin-Streptomycin (Gibco 15140122). Mouse Embryonic Stem Cells (mESCs) were cultured using standard culture procedures. These cells were isolated by Jerry Crabtree’s laboratory (Stanford University). Cells were maintained feeder-free using Knock-Out DMEM (Gibco 10829018) media containing, 7.5% Knock-out Replacement Serum (Gibco 10828028), 7.5% ES-sure FBS (Applied StemCell ASM-5017), 1% Penicillin-Streptomycin (Gibco 15140122), 1% Glutamax (Gibco 35050061), 1% MEM Non-Essential Amino Acids (Gibco 11140050), 1% Sodium Pyruvate (Gibco 11360070). LIF was replaced daily, and ES cells were passaged every 48 hours.

### Lentivirus Transduction

HEK 293T cells were transfected with lentiviral constructs and packaging plasmids (psPAX2 and pMD2.G) using PEI transfection (Polysciences). Two days post-transfection, the virus-containing cell culture media was collected, filtered with a 0.44 μm syringe filter, and centrifuged at 50,000 g for 2.5 hours at 4°C (SW28 rotor on ultracentrifuge). The viral pellet was resuspended in PBS and used for subsequent infections. Selection of lentiviral constructs was achieved with the following doses: puromycin (2 μg/mL), blasticidin (10 μg/mL), hygromycin (150 μg/mL), and zeocin (200 μg/mL).

### Antibodies used in this study

rabbit polyclonal antibody ɑ-PHF10 (Invitrogen PA5-30678); Co-IP, ChIP, WB

rabbit ɑ-ARID2 (Invitrogen PA5-35857); Co-IP, WB

mouse ɑ-V5 Tag (E9H8O) (Cell signaling technology #80076); Co-IP, WB

rabbit ɑ-V5 Tag (D3H8Q)(Cell Signaling Technology #1302); ChIP, WB

rabbit ɑ-ARID1A (D2A8U)(Cell Signaling Technology #12354); ChIP

rabbit ɑ-H3K27me3 (C36B11)(Cell Signaling Technology #9733); ChIP

rabbit ɑ-SS18 (D6I4Z)(Cell Signaling Technology #21792); ChIP

rabbit polyclonal ɑ-BAF60A (Bethyl Laboratories #A301-595A);

ChIP rabbit polyclonal ɑ-H3 c-terminal (Epicypher #13-0001); ChIP

rabbit ɑ-H2AK119ub (D27C4)(Cell Signaling Technology #8240); ChIP

rabbit ɑ-SMARCC1/BAF155 (D7F8S) (Cell signaling technology #11956S); WB

mouse ɑ-BRG1 (H-10) (Santa Cruz Biotechnology sc-374197); WB

rabbit polyclonal ɑ-BAF57 (Bethyl Laboratories A300-810A); ChIP

rabbit recombinant ɑ-Histone H3.3 (Active Motif 91191); ChIP

rabbit ɑ-Histone H4 (D2X4V) (Cell signaling technology #14149H4); ChIP

goat anti-mouse IgG IRDye 680RD polyclonal secondary antibody (Li-Cor)

goat anti-mouse IgG IRDye 800CW polyclonal secondary antibody (Li-Cor)

goat anti-rabbit IgG IRDye 680RD polyclonal secondary antibody (Li-Cor)

goat anti-rabbit IgG IRDye 800CW polyclonal secondary antibody (Li-Cor)

### Ammonium Sulfate Protein Extraction

Cells were plated onto 15 cm^2^ plates, and 30 × 10^6^ cells were harvested after 48 hours. After a single wash with PBS to remove any remaining media, the cells were lysed in 10 mL of the lysis buffer (10 mM HEPES pH 7.5, 25 mM KCl, 1 mM EDTA, 0.1% NP-40, 10% glycerol, 1 mM DTT, protease inhibitor tablet (Roche), 1 mM sodium orthovanadate, and 10 mM sodium butyrate) and incubated for 10 minutes on ice. Washing with lysis buffer one more time, lysed cells were resuspended into 660 μL of the resuspension buffer (10 mM HEPES pH 7.5, 100 mM KCl, 1 mM EDTA, 3 mM MgCl_2_, 10% glycerol, 1 mM DTT, protease inhibitors cocktail (10 μg/mL of Leupeptin, Chymostatin, Pepstatin A), 1 mM sodium orthovanadate, 10 mM sodium butyrate and 300 mM ammonium sulfate) and incubated for 30 minutes at 4°C. After moving into the centrifugation tube (Beckman Coulter polycarbonate tube #343778), it is spun down with ultracentrifugation (ThermoFisher rotor S140-AT) at 100,000 rpm for 10 minutes. The supernatant was incubated with 200 mg solid ammonium sulfate for 45 minutes at 4°C and spun down again with ultracentrifugation at 100,000 rpm for 10 minutes. The pellets are frozen and stored at −80°C.

### Co-Immunoprecipitation

Protein pellets were resuspended in 220 μL of IP buffer (20 mM HEPES pH 7.5, 150 mM KCl, 1 mM EDTA, 1 mM MgCl_2_, 10% glycerol, 0.1% Triton X-100, 1 mM DTT, protease inhibitors cocktail, 0.2 mM sodium orthovanadate, 10 mM sodium butyrate). Protein G beads washed with PBS are incubated with each antibody, mouse ɑ-V5 Tag (E9H8O) (1:50, Cell signaling technology), rabbit ɑ-PHF10 (1:100, Invitrogen), and rabbit ɑ-ARID2 (1:100, Invitrogen) in PBS for 1 hour at room temperature. After incubation, protein G beads are washed with PBS two times and washed with an IP buffer two times. Protein G beads were incubated with 400 μL of protein extracts with the protein concentration of 0.75 μg/μL overnight at 4°C. Supernatants were saved as a flow-through, and the beads were washed with an IP buffer five times on ice. IPs were extracted with 50 μL of RIPA buffer (50 mM Tris pH 7.8, 150 mM NaCl, 1% NP-40, 1% Sodium dodecyl sulfate (SDS), 0.1% Sodium deoxycholate (DOC), 1 mM DTT, and protease inhibitor cocktails) by boiling for 5 minutes.

### Chromatin Immunoprecipitation

ChIP samples were prepared by fixing one of two ways and then processed the same after the fixation steps.

#### Single Formaldehyde Fixation

Cells were harvested by first washing with 10 mL PBS (Fisher Scientific), then were incubated in trypsin-EDTA 0.25% (Fisher Scientific) for 5 minutes. Cells were washed with PBS and were fixed using a final concentration of 1% formaldehyde for 10 minutes. The addition of 0.125% glycine then quenched fixation. This method was used for all samples except the ones listed below.

#### Double Fixation method

Detailed methods (an adaptation of Bing and Brasier^26^) are available at dx.doi.org/10.17504/protocols.io.yxmvm396nl3p/v1. This fixation method was used to map the enrichment of other BAF subunits at the recruitment locus (Supplemental Figure 2).

#### Preparation of Chromatin and IP

Cell pellets (30 × 10^6^ cells) were resuspended in CiA NP-Rinse 1 (50 mM HEPES pH 8.0, 140 mM NaCl, 1 mM EDTA, 10% glycerol, 0.5% NP-40, 0.25% Triton X-100) and incubated on ice for 10 minutes before being spun down at 1000 g for 5 minutes at 4°C. The supernatant was aspirated, and cells were resuspended in CiA NP-Rinse 2 (10 mM Tris pH 8.0, 1 mM EDTA, 0.5 mM EGTA, 200 mM NaCl) and spun down at 1000 g for 5 minutes at 4°C. The supernatant was aspirated, and the salt was washed from the sides of the tube by gently adding 5 mL of Covaris Shearing Buffer (0.1% SDS, 1 mM EDTA pH 8.0, 10 mM Tris HCl pH 8.0) along the ridge of the tube while rotating clockwise so that the buffer trickled down the sides and did not disturb the pellet. The samples were spun down at 1000 g for 3 minutes at 4°C, and the supernatant was carefully aspirated. This step was repeated one additional time before the pellet was resuspended in 0.9 mL of Cia Covaris Shearing Buffer (0.1% SDS, 1 mM EDTA pH 8.0, 10 mM Tris HCl pH 8.0) + Protease Inhibitor cocktail (1:1000) and transferred to a Covaris glass tube.

### Sonication

Cells were sonicated for 8 minutes (single fixation ChIP) or 10 minutes (double fixation ChIP) to generate DNA fragments of the desired size using a Covaris E220 Evolution at 5.0 duty factor, 140 peak power, and 200 cycles per burst. Following this, samples were transferred to microcentrifuge tubes and spun at 10,000 g for 5 minutes at 4°C, and the supernatant (chromatin stock) was transferred to a new tube. Then, 25 μL of chromatin stock was aliquoted as input DNA. All sonicated chromatin was stored at −20 °C until ready for IP.

### Immunoprecipitation

Sonicated chromatin was diluted in 5x IP buffer (250 mM HEPES, 1.5 M NaCl, 5 mM EDTA pH 8.0, 5% Triton X-100, 0.5% DOC, and 0.5% SDS) to a concentration of 1x IP Buffer and incubated overnight with 10 μL Pierce protein A/G beads (Fisher Scientific) at 4°C with rotation. Beads were collected on a magnet and washed with 1x IP buffer for 3-5 minutes with rocking. Beads were then washed with 0.5 mL DOC buffer (10 mM Tris pH 8, 0.25 M LiCl, 0.5% NP-40, 0.5% DOC, 1 mM EDTA) for 1-3 minutes with rocking and then were washed with 1 M TE pH 8.0. The residual supernatant was removed.

### Reverse Cross-Linking

Beads incubated as part of IP were resuspended and mixed well in 100 μL TE, 5 μL 10% SDS, and 5 μL Proteinase K (10 mg/mL). Inputs were resuspended and mixed well in 75 μL TE, 5 μL 10% SDS, and 5 μL Proteinase K (10 mg/mL). Samples were then incubated at 55°C for 3 hours, followed by an incubation at 65°C overnight.

### Elution

Supernatant was collected from each sample and transferred to a new tube. Then, 550 μL of NTB Binding Buffer (Machery Nagel) was added to each sample, and samples were loaded into NucleoSpin PCR/gel clean-up columns (Machery Nagel). Columns were washed with 750 μL of Buffer PE (80% EtOH, 10 mM tris pH 7.5). IPs were eluted with 25 μL of Buffer EB (Machery Nagel), and inputs were eluted with 50 μL of Buffer EB.

*ChIP-qPCR:* pPCR samples were prepared using Accuris qMAX SYBR Green (MidSci), according to the manufacturer’s instructions. Analysis of qPCR samples was performed on a QuantStudio 3 Flex system (Life Technologies). For ChIP-qPCR experiments, enrichment (bound over input) values were normalized to values with no RAP treatment (Rap/No Rap) and then to a control locus (*HoxA9*) enriched for polycomb-associated histone marks.

### Transcriptional Analysis

Cells were plated onto 6 cm^2^ plates and left to attach overnight. For FIRE-dCas9 lines, 3 nM Rapamycin was added to experimental plates, and vehicle was added to control plates 24 hours before RNA isolation using Trisure (Bioline) per the manufacturer’s instructions. Before Trisure treatment, cells were visualized under the microscope to ensure subconfluency and stem cell morphology. cDNA was made from 1 μg of total RNA using the Superscript VIVO mix (Invitrogen). cDNA was diluted 1:4 and 2 μL was used in a 20 μL qPCR Taqman assay (Invitrogen). Assay probes were used to detect cDNA from *Nkx2.9* (Mm00435145_m1). *Ywhaz* (Mm01722325_m1) was used for normalization^27^. An mESC cell line constitutively expressing dCas9-VP64^20^ with the same sgRNAs to *Nkx2.9* used in the FIRE-dCas9 experiments was used as a positive control.

### Western Blots

Whole cell extract of each sample was prepared using RIPA Buffer (1% SDS), Protease Inhibitor Cocktail (1:1000), 1 M DTT (1:1000), and Benzonase nuclease (Sigma Aldrich) (1:200). Proteins were then separated by SDS-PAGE electrophoresis with a 4-12% Bis-Tris protein gel (Fisher Scientific) in 1x MOPS SDS Running Buffer (Fisher Scientific). Bands were transferred to an Immobilon-FL PVDF membrane (Fisher Scientific), then blocked in Intercept Protein-Free Blocking Buffer (Li-Cor) for one hour. The membrane was then incubated overnight with Primary Antibodies diluted (1:1000) in Intercept T20 Antibody Diluent (Li-Cor).

The membrane was washed four times in TBS-T (0.2% Tween 20) for 5 minutes in each wash and probed with IRDye fluorescence-conjugated secondary antibody (Li-Cor) that was diluted (1:20,000) in T20 Antibody Diluent (Li-Cor) with 0.01% SDS. After, the membrane was washed four times in TBS-T (0.2% Tween 20) before a final rinse in TBS to remove the residual Tween 20. Bands were detected using an Odyssey DLx imaging system (Li-Cor). Raw western blots used in figures are provided in Supplemental Figure 3.

### Plasmids used or generated in this study

All plasmids used in this study were confirmed by nanopore-based whole plasmid sequencing (Plasmidsaurus or OMRF Clinical Genomics Center). The generated plasmids will be submitted to Addgene upon peer-reviewed publication of this manuscript but are immediately available by contacting the corresponding author.

### MS2-stem-loop sgRNAs targeting *Nkx2.9* were previously described in Braun and Kirkland et al.,^15^ Supplementary Table 1

sgRNA1: GGGGCGGGTGCCGGGCGGGG

sgRNA2: GGGGCGGAGATGGCACCTTC

sgRNA3: ACCAAAGTGGGGACAGATAA

dCas9: Lv EF1a dCas9-HA 2A Blast was a gift from Jerry Crabtree (Addgene plasmid # 102812 ; http://n2t.net/addgene:102812 ; RRID:Addgene_102812)

MS2-2xFrb (Hygro^R^): pJK192 generated for this study

PHF10-2xFkbp-V5 (Puro^R^): pJK314 generated for this study

DPF2-2xFkbp-V5 (Puro^R^): pJK315 generated for this study

BRD9-2xFkbp-V5 (Puro^R^): pJK343 generated for this study

lenti dCAS-VP64_Blast was a gift from Feng Zhang (Addgene plasmid # 61425 ; http://n2t.net/addgene:61425; RRID:Addgene_61425)

psPAX2 was a gift from Didier Trono (Addgene plasmid # 12260 ; http://n2t.net/addgene:12260 ; RRID:Addgene_12260)

pMD2.G was a gift from Didier Trono (Addgene plasmid # 12259 ; http://n2t.net/addgene:12259 ; RRID:Addgene_12259)

### Primers used in this study

JK800 Nkx2.9 (−2455) AAA TGA CCG GGC TCT GTA TG

JK801 Nkx2.9 (−2455) AGT TCC CGC TTC ACA TTC TC

JK429 mNkx2.9 (−820) CTC CAT TCG AGG ACC CAA GG

JK430 mNkx2.9 (−820) CTG CTA ACT GGC ACC GAC TT

JK431 mNkx2.9 (−315) TCT TGG GTG GCG AAC AGT G

JK432 mNkx2.9 (−315) AAT AAA GTC GCT CCA CCC TCC

JK433 mNkx2.9 (+160) CCG CTC CTA AGG ATG GAA GT

JK434 mNkx2.9 (+160) TTC AAA GCC CTC CGA GTA GC

JK435 mNkx2.9 (+630) ATC CCG GTC TTT TCG GAT CG

JK436 mNkx2.9 (+630) TGC GTC TGA GTC CAC ACA TC

JK437 mNkx2.9 (+1400) ACC TCT GCC GTT GTT GCT C

JK438 mNkx2.9 (+1400) GCC TTC GGA TAT GGC AGC AT

JK342 mGapdh F ChIP CTC TGC TCC TCC CTG TTC C

JK343 mGapdh R ChIP TCC CTA GAC CCG TAC AGT GC

JK344 mHoxa9 F ChIP AAG AAG GAA AAG GGG AAT GG

JK345 mHoxa9 R ChIP TCA CCT CGC CTA GTT TCT GG

## Acknowledgments

This work was supported in part by grants from NIH/NIGMS 5P20 GM103636 (Research Project Leader: JGK), the Stephen M. Prescott Endowment Fund for the Best and Brightest (JGK), and an OMRF pre-doctoral scholarship (JYP).

---

**Supplemental Figure 1:**
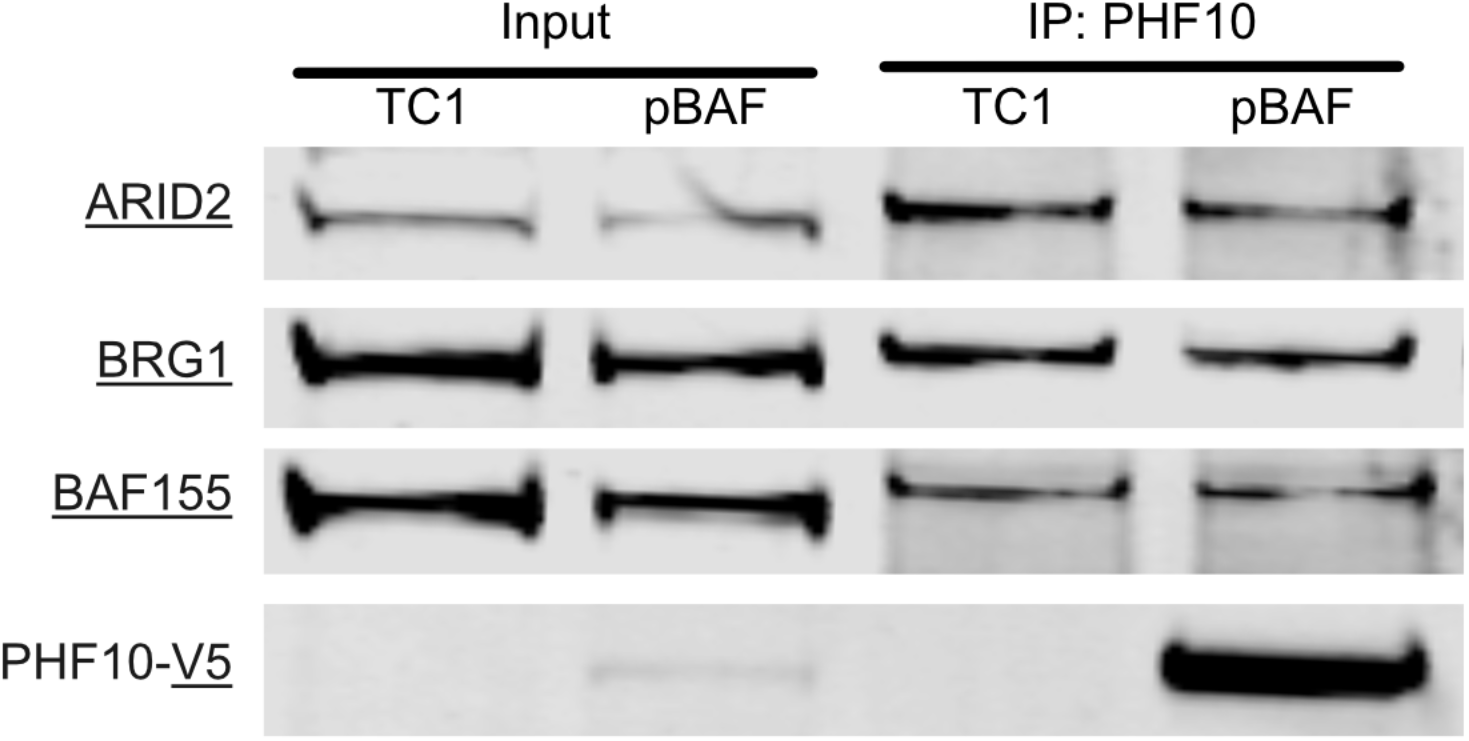
Co-IP pulldown with PHF10. Representative western blot from a co-IP. Input and pulldown with an antibody against PHF10. PHF10 interacts with ARID2, BRG1, and BAF155. The PHF10 pulldown also pulls down the PHF10-2xFkbp-V5 version of the protein expressed in our pBAF FIRE-Cas9 cells, but not in wild-type TC1 cells as detected by an antibody against the V5 epitope.

**Supplemental Figure 2:**
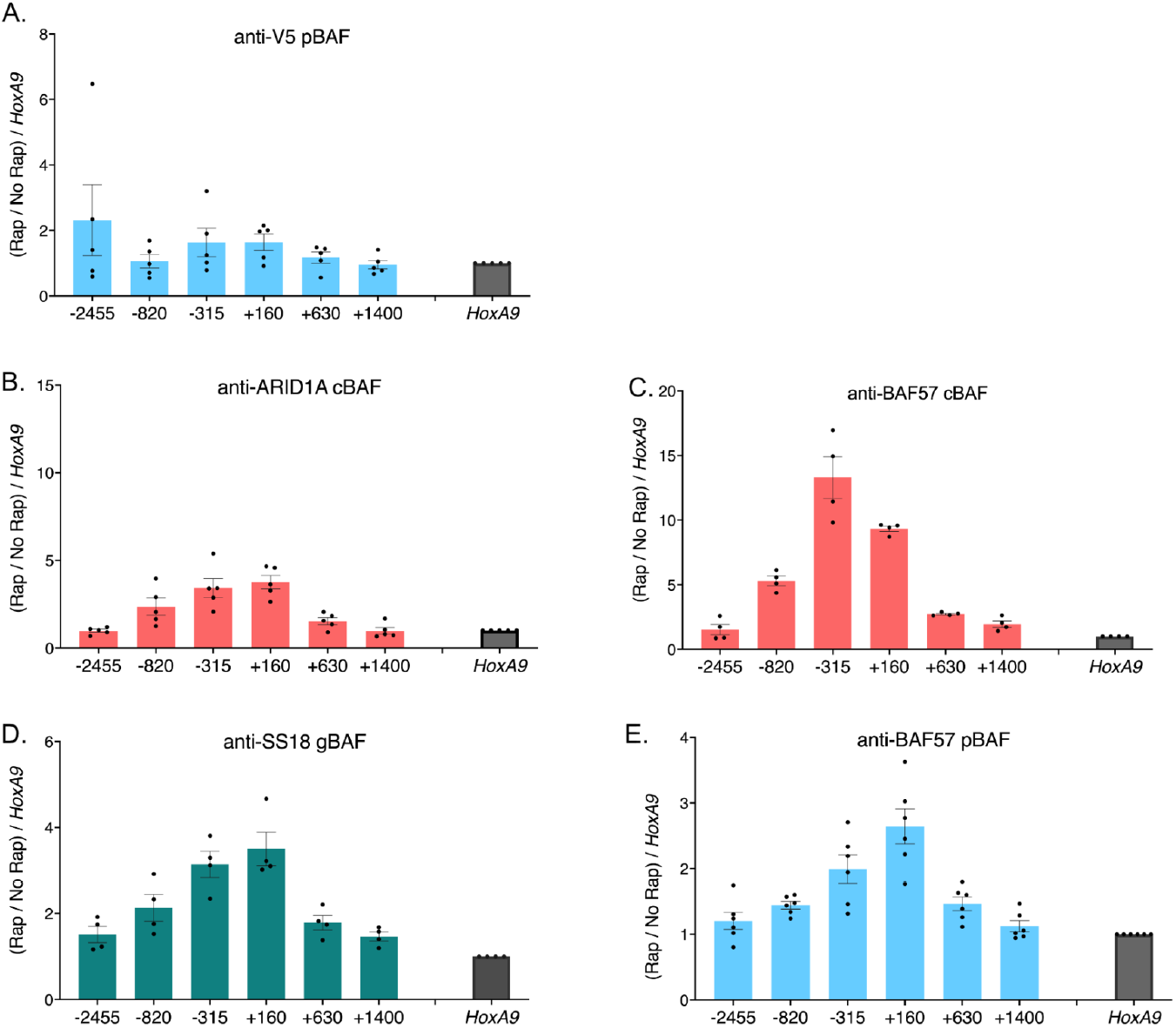
Double-fixation ChlPs. Chromatin immunoprecipitations against various antibodies to determine enrichment for other BAF subunits during recruitment at the *Nkx2.9* locus. **A**. Lack of enrichment with V5 pulldowns of PHF10-2xFkbp-V5 proteins **B**. ARID1A enrichment with cBAF recruitments **C**. BAF57 enrichment with cBAF recruitments **D**. SS18 enrichment with gBAF recruitments **E**. BAF57 enrichment with pBAF recruitments. Data is presented as Rap/No Rap (recruitment/no recruitment) normalized to the same conditions at the *HoxA9* locus where no BAF complex is being recruited (n=4-6; s.e.m.)

**Supplemental Figure 3:**
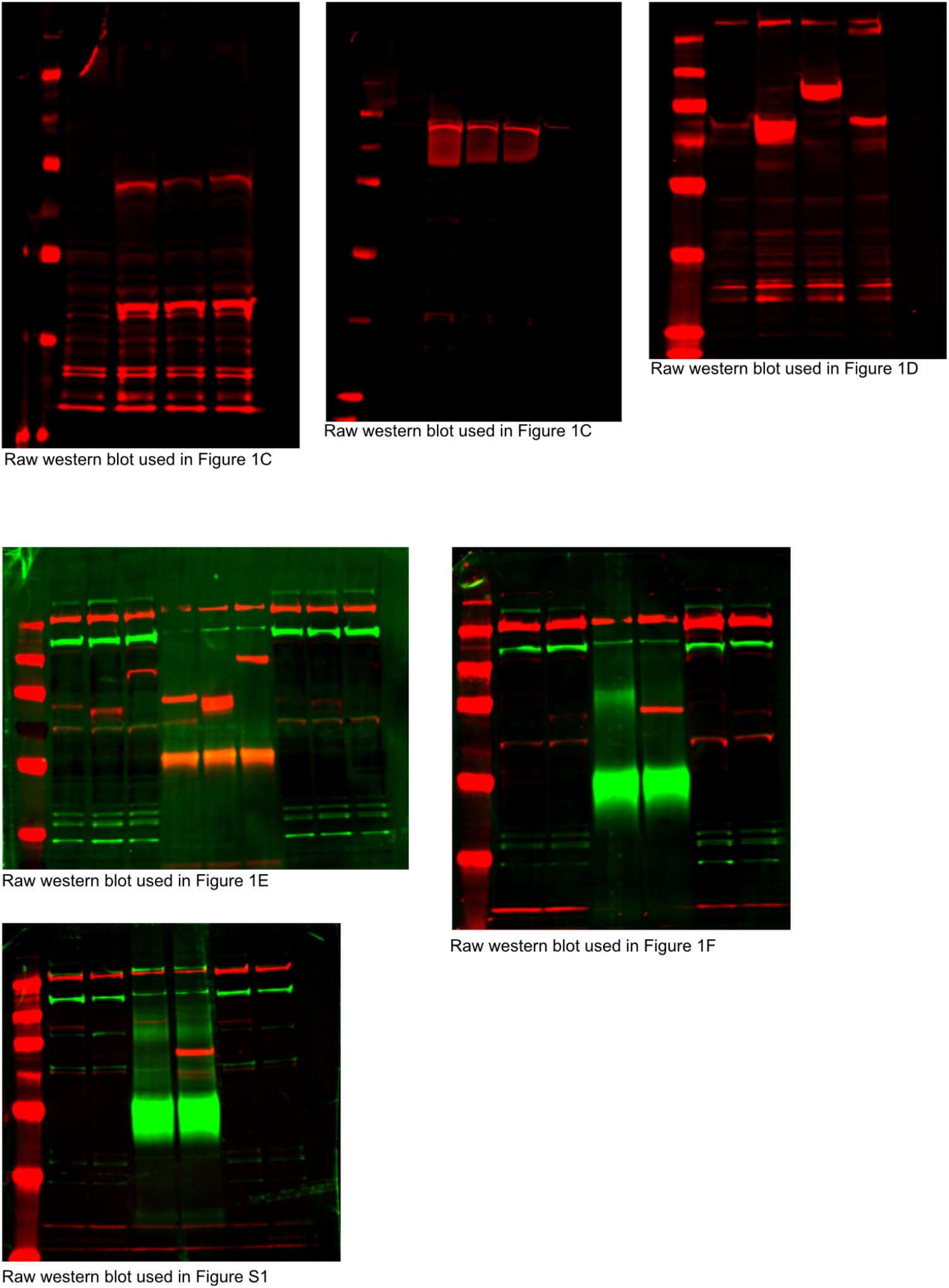
Raw western blots used in figures.

**Supplemental Figure 4:**
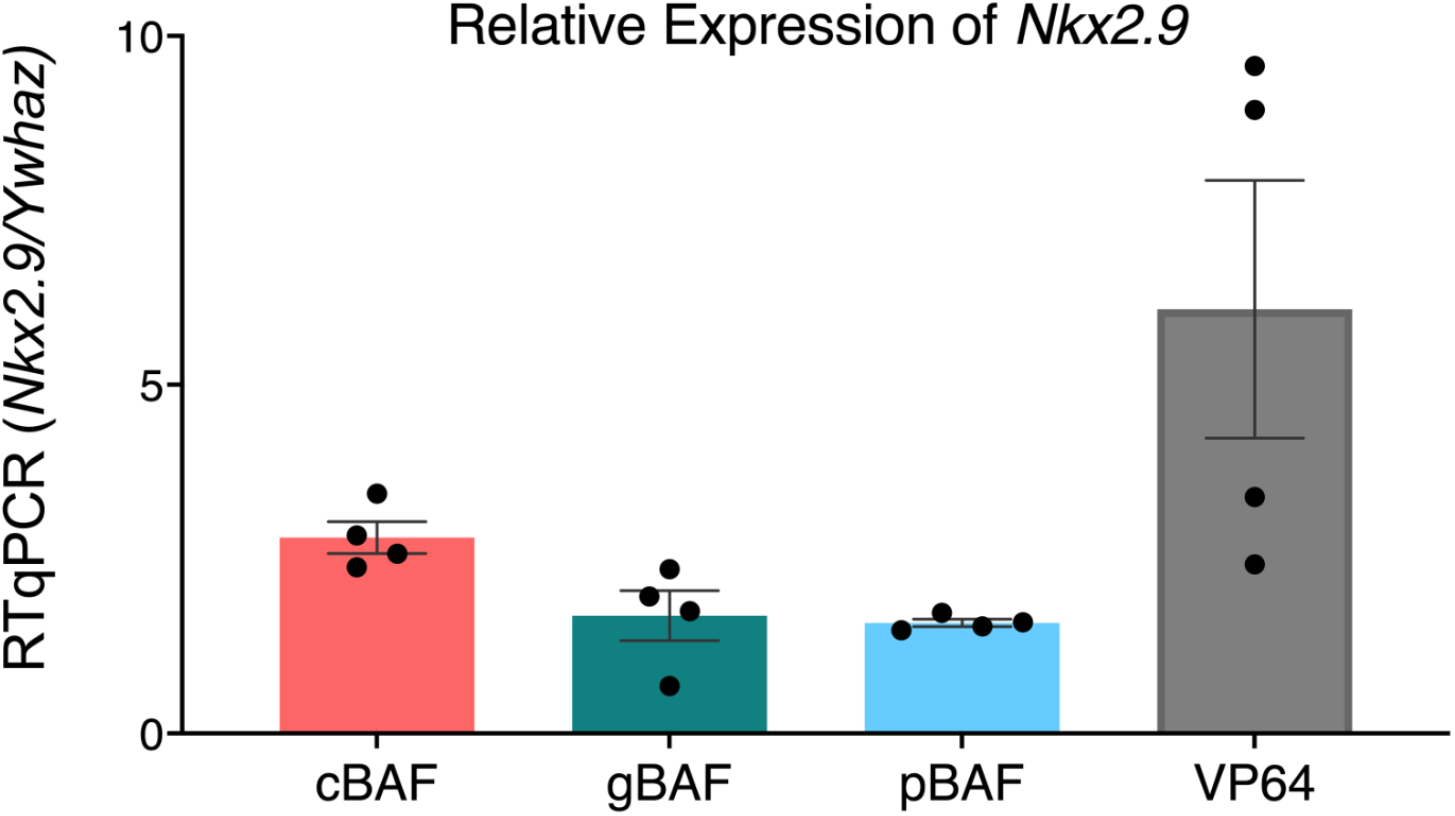
Relative expression of NKX2.9 including a dCas9-VP64 positive control. For the BAF complexes data is presented as 2^(Ct *Ywhaz*-Ct *Nkx2.9*) with a fold change of Rap/No Rap (recruitment/no recruitment). For the CRISPR-A data is presented as 2^(Ct *Ywhaz-* Ct *Nkx2.9*) with a fold change of sgRNAs/no sgRNAs (recruitmenUno recruitment). (n=4; s.e.m.)

